# DCBK: A Fragment-Based Hybrid Simulation and Machine Learning Framework for Predicting Ligand Dissociation Kinetics

**DOI:** 10.1101/2025.11.01.685950

**Authors:** Yang Li, Yixin Wang, Xueying Zheng, Jinjiang Guo, Rumin Zhang

## Abstract

Predicting small-molecule dissociation kinetics remains challenging due to the high computational cost of simulating rare unbinding events and the limited scalability of existing approaches. Here, we introduce DCBK (Divide-and-Compute Binding Kinetics), a physics-informed, fragment-resolved framework for predicting ligand dissociation across diverse protein–ligand systems. DCBK integrates steered molecular dynamics and umbrella sampling to reconstruct unbinding pathways and free energy profiles, followed by BRICS-based ligand fragmentation. Fragment-level energetic and structural features are combined with machine learning trained on experimental kinetics data, enabling efficient prediction of dissociation rate constant (k_off_) values. Validation across five protein families demonstrates strong agreement with experiments, with ablation and unseen-target analyses highlighting the contribution of fragment-level features and the framework’s transferability. Importantly, DCBK allows dissociation kinetics to be interpreted in terms of fragment-level contributions, offering mechanistic insight and guiding ligand modification to optimize residence time. This interpretable and extensible framework links molecular structure, energetics, and kinetics, providing actionable insights for lead optimization and rational drug design.

## Introduction

Understanding ligand–target interactions through binding kinetics is increasingly recognized as a key component of rational drug discovery, particularly in lead optimization[1, 2]. While equilibrium binding affinity quantifies interaction strength, kinetic parameters such as the association rate constant (k_on_) and dissociation rate constant (k_off_) provide mechanistic insight into the speed and lifetime of target engagement, closely linked to pharmacodynamic efficacy and selectivity[3]. Compounds with similar affinities can display markedly different residence times, resulting in significant variations in therapeutic outcomes. For instance, ligands targeting HSP90 show comparable affinities but up to ninefold differences in residence time, whereas CDK8/CycC inhibitors exhibit extended residence times from minutes to hundreds of minutes, attributable to strengthened hinge-region interactions—demonstrating that optimizing affinity alone is insufficient and highlighting the importance of quantitative modeling of dissociation kinetics.[4, 5]. These cases underscore that optimizing binding affinity alone is insufficient, and that quantitative modeling of binding kinetics—particularly residence time, which is inversely related to the k_off_—is critical for designing drugs with durable efficacy. This drives the need for computational methods capable of capturing both equilibrium thermodynamics and atomic-level kinetic barriers.

Experimental in vitro techniques measuring binding kinetics spanning a vast timescale of eight orders of magnitude from sub-seconds to hours and days, including time-resolved fluorescence, stopped- or quenched flow, biosensors such as surface plasmon resonance (SPR), global progress curve analysis (GPCA), radioligand or fluorescent ligand binding, and jump dilution, enable precise measurement of protein–ligand association and dissociation rates, generating valuable kinetic data that inform mechanistic understanding of ligand–target interactions[6, 7]. Complementing these experimental resources, accurate computational prediction of k_off_ can prioritize compounds with favorable kinetic profiles at early stages of drug discovery, accelerating the identification of candidates with durable efficacy and optimal pharmacological properties.

However, direct computation of binding kinetics remains computationally demanding and requires extensive knowledge of the system and careful molecular dynamics (MD) sampling strategies[8]. Thermodynamically, the binding free energy (ΔG_bind_) reflects the stability difference between bound and unbound states, whereas kinetics depend on the entire free energy landscape connecting these states through a dissociation transition state. The activation free energy barrier ( Δ*G*^‡^) is typically high, meaning that spontaneous unbinding events are rarely observed within conventional simulation timescales; for instance, a 100 ns unbiased MD trajectory is unlikely to capture dissociation events when ligand residence times extend to hours or longer. While landmark multi-microsecond simulations from D. E. Shaw Research have demonstrated that unbiased MD can reveal dissociation mechanisms in select systems, such studies remain computationally intensive.. Likewise, ligand-GaMD and other enhanced sampling techniques have been successfully used to investigate unbinding pathways and estimate dissociation rates[9, 10], but their application is often constrained by system size and pathway complexity.

Numerous studies have demonstrated that a widely adopted and effective strategy for computing ligand dissociation kinetics is a two-stage approach. The first stage employs enhanced sampling or accelerated dynamics techniques to efficiently explore unbinding pathways and generate candidate reaction coordinates. Representative techniques include τ-Random Acceleration MD (τ-RAMD), Steered MD (SMD), Adaptive Steered MD (ASMD), Metadynamics (MetaD), Scaled MD, and Adaptive Biased MD (ABMD) [11] etc., which bias the system or modify the potential energy landscape to facilitate ligand dissociation. Complementary approaches, such as Site Identification by Ligand Competitive Saturation (SILCS), can further inform the selection of plausible dissociation pathways. Building on the pathways identified in the first stage, the second stage involves rigorous free energy reconstruction, typically performed using methods such as Adaptive Steered MD (ASMD) combined with Jarzynski averaging or MetaD-derived free energy surface analysis. More computationally intensive techniques, including US, nudged elastic band (NEB) calculations[12], and attach–pull–release (APR) protocols[13], offer highly detailed characterization of the dissociation transition state. These physics-based approaches reconstruct the free energy surface along a chosen reaction coordinate and estimate the Δ*G*^‡^, which can then be related to k_off_ through transition state theory. While these physics-based reconstructions are rigorous and mechanistically interpretable, their substantial computational cost and complex setup (e.g., reaction coordinate selection, biasing windows, and restraint schemes) render them impractical for high-throughput screening.

In parallel, data-driven approaches have been developed to directly predict k_off_ from molecular structure. Quantitative structure–kinetics relationship (QSKR) models and ML frameworks, including Random Forest Regressor (RF), Gradient Boosting, and Graph neural Networks (GNNs), have been applied to experimental kinetic datasets to capture complex structure–kinetic relationships. Publicly available resources such as the KIND dataset[14] (3,812 small molecules with measured k_on_, k_off_, and K_d_), the *PDBbind-koff 2020* compilation[15] (∼680 protein–ligand complexes), and the 501-entry unbinding rate dataset curated by Amangeldiuly et al[16]. provide valuable training and benchmarking material, although they remain limited relative to traditional affinity datasets and cover restricted chemical and target space. Building on these datasets, several predictive models have been developed[17, 18], including ModBind, exemplifying how ML can be applied to estimate k_off_. Collectively, these studies demonstrate that while ML can efficiently predict ligand dissociation rates and aid in ligand optimization, purely data-driven approaches still face challenges in generalization and mechanistic interpretability, limiting their ability to provide detailed molecular insight into the determinants of binding kinetics.

Building upon these developments, we developed DCBK (Divide and Compute Binding Kinetics), a hybrid framework integrating atomistic simulations and ML to achieve accurate and interpretable k_off_ predictions. DCBK first explores unbinding pathways via SMD, reconstructs free energy barriers with umbrella sampling, and decomposes ligands into BRICS-based fragments to quantify fragment-level energetic contributions. These features are subsequently refined using a data-driven correction model. To systematically evaluate the approach, we constructed a mechanistically annotated kinetic database comprising 60 ligand–protein complexes across five protein families. This benchmark integrates atomistic dissociation pathways, computed activation free energies, and experimental k_off_ values, providing a scalable foundation for predictive modeling and mechanistic understanding of ligand dissociation kinetics. Importantly, DCBK enables interpretation of dissociation kinetics at the fragment level, offering actionable insights for guiding ligand modification and optimizing residence time.

## Results

### Overview of the Hybrid Workflow

Ligand dissociation from protein targets is a key determinant of drug efficacy, with residence time often more predictive of in vivo activity than equilibrium binding affinity. To capture the mechanistic basis of dissociation kinetics and enable accurate predictions, we developed DCBK (Divide and Compute Binding Kinetics), a hybrid computational workflow that integrates physics-based enhanced sampling with machine learning (Fig. 1). The energetic rationale for this framework is summarized in Fig. 1A, which illustrates how ligands with similar overall binding free energies can still display different dissociation rates when their unbinding barriers differ. The workflow begins with a curated protein-ligand complex, from which both intact ligands and chemically meaningful fragments are incorporated into the simulation system. Dissociation pathways are generated through supervised SMD, providing physically interpretable trajectories that inform subsequent free-energy reconstruction. Umbrella sampling is then applied along the dissociation coordinate to compute ligand-level activation barriers (whole-ΔG^‡^), quantifying the energetic difficulty of unbinding and establishing a foundation for fragment-level analysis.

**Figure 1.**
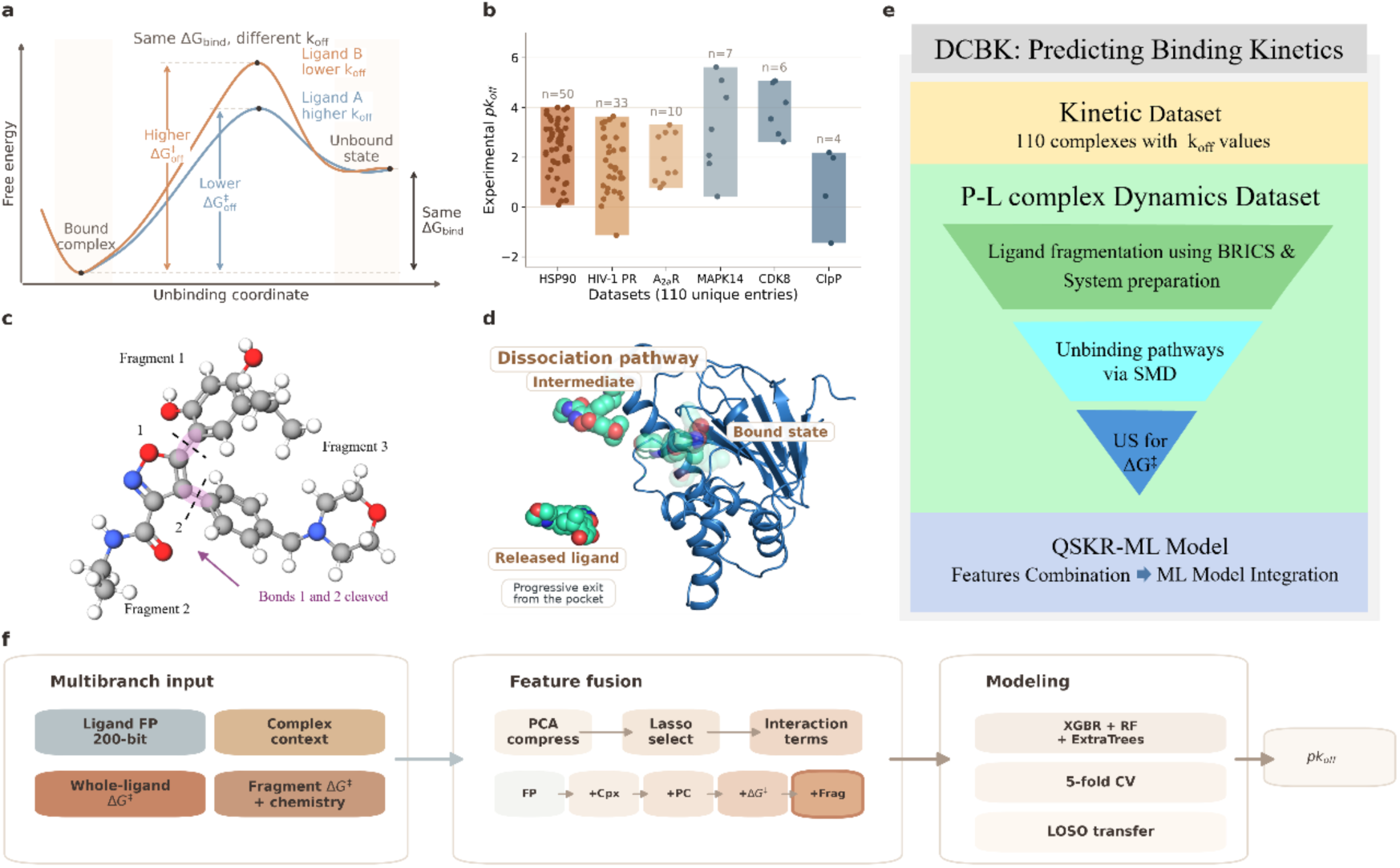
Overview of the hybrid DCBK workflow for predicting ligand dissociation kinetics. (A) Conceptual free-energy profiles show that ligands with similar overall ΔG_bind_ can still exhibit different k_off_ because their unbinding barriers differ. (B) Distribution of experimental pk_off_ values across the six curated protein systems in the study (HSP90, HIV-1 PR, A2aR, MAPK14, CDK8, and ClpP), spanning 110 unique entries and a broad kinetic range. (C) Example of BRICS fragmentation applied to a ligand, highlighting chemically meaningful substructures retained for downstream simulation and analysis. (D) Representative ligand dissociation pathway obtained from SMD simulations, with the protein shown in cartoon representation and ligand positions fading from the bound state to the released state along the unbinding route. (E) Overall DCBK workflow linking the kinetic dataset and protein-ligand dynamics dataset to fragmentation, pathway generation, free-energy reconstruction, and ML integration. (F) Compact schematic of the DCBK-ML model, summarizing the multibranch inputs, feature-fusion steps, ensemble modeling stage built from XGBR, RF, and ExtraTrees, and the final pk_off_ prediction.

To characterize the kinetic diversity of the systems, we curated experimental dissociation rates (k_off_) for six protein systems, encompassing 110 unique protein-ligand entries: heat shock protein 90 (HSP90), adenosine A_2A_ receptor (A_2A_R), cyclin-dependent kinase 8 (CDK8), mitogen-activated protein kinase 14 (MAPK14), human immunodeficiency virus type 1 protease (HIV-1 protease), and caseinolytic protease complex ClpP_1_P_2_ (ClpP). The measured −log_10_*k*_off_(pk_off_) values span a broad range from approximately −2 to 6, covering both transiently bound ligands and slow-dissociating inhibitors (Fig. 1B, Table S1). Each entry includes the experimentally measured kinetics together with the corresponding protein-ligand structural model, providing a unified dataset of kinetic and structural information. This dataset serves as the basis for subsequent mechanistic simulations at both the ligand and fragment levels, as well as for training the DCBK machine-learning model. For each complex, both whole-ligand and fragment-level MD trajectories were generated to capture complete unbinding transitions, enabling computation of ligand-level and fragment-level activation barriers 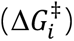 that are then integrated into the predictive ML framework.

Each ligand was decomposed into chemically meaningful fragments using the BRICS scheme (Fig. 1C), which preserves core ring systems while segmenting flexible linker regions. For each fragment, supervised SMD was performed to generate atomistic unbinding trajectories, capturing detailed motions of individual atoms as the ligand progresses from the bound to dissociated state (Fig. 1D). These SMD-derived pathways, combined with BRICS-based fragment decomposition, enabled computation of fragment-level activation barriers, providing a physically interpretable map of which subunits govern residence time and highlighting potential hotspots for rational design. Along these pathways, US was applied to rigorously reconstruct free-energy profiles for both the whole ligand and its constituent fragments. Importantly, ligands with similar overall binding free energies (ΔG_bind_) can still exhibit substantially different dissociation kinetics because fragment-specific barriers reshape the unbinding landscape, demonstrating why binding affinity alone is insufficient to predict k_off_ (Fig. 1A). This integrated fragment-resolved energetic analysis establishes a mechanistic basis for understanding experimentally observed differences in residence time and provides essential input for subsequent machine-learning modeling.

Fragment- and ligand-level features, including computed activation barriers, molecular descriptors (for example RDKit-derived logP, TPSA, molecular weight, and rotatable-bond counts), chemical structure information, and protein-ligand interaction features such as hydrogen-bond counts, contact frequencies, and pocket fingerprints, were integrated into a machine learning model denoted DCBK-ML, which builds upon the QSKR framework (Fig. 1F). In the revised compact schematic, the ensemble modeling stage is explicitly represented by Extreme Gradient Boosting Regressor (XGBR), RF, and Extremely Randomized Trees Regressor (ExtraTrees). By capturing nonlinear correlations and systematic deviations not fully represented by simulations, such as slow conformational rearrangements and solvent-mediated effects, DCBK-ML refines predictions of experimental pk_off_ while preserving fragment-level interpretability. This hybrid approach therefore combines mechanistic insight from enhanced-sampling simulations with data-driven generalization across diverse chemical scaffolds.

Building upon these features, we further integrated the complete set of whole-ligand and fragment-level MD simulation trajectories with corresponding experimental *k*_off_measurements into a structured database. This compilation captures full unbinding transitions, computed free-energy barriers, and ligand–protein interaction details, providing a mechanistic and quantitative foundation for analysis. The resulting unified platform enables rigorous statistical assessment and machine-learning modeling, directly linking atomistic dynamics with predictive, data-driven interpretations of ligand dissociation kinetics.

### Validation Across Diverse Systems

To evaluate the predictive generality of DCBK-ML, we compiled a curated dataset of 110 protein–ligand complexes spanning five distinct target families: HSP90 (n = 50), CDK8 (n = 6), HIV-1 protease (n = 33), ClpP (n = 4), p38α MAPK (n = 7), and A_2A_R (n = 10). These targets encompass human, bacterial, and viral proteins, exhibiting diverse structural folds, binding-pocket architectures, and molecular recognition mechanisms. Experimental dissociation rates (p*k*_off_) ranged from approximately −2 to 6, covering nearly eight orders of magnitude and providing a stringent test for both rapid and slow dissociating ligands. For each complex, both intact-ligand and fragment-level trajectories were generated to capture complete unbinding transitions using SMD. The pulling coordinate was defined as the distance between the ligand and the protein binding pocket, specifically calculated using the three closest protein–ligand atomic pairs, to ensure a physically meaningful representation of the dissociation process (see Methods). Throughout the simulations, we monitored ligand–pocket contact numbers and solvent-accessible surface area to confirm pathway integrity and avoid artificial steric crossings. Unreasonable SMD pathways were discarded or rerun to ensure reliability. On these validated pathways, US simulations were carried out to reconstruct the free energy profiles and *ΔG*^‡^. All resulting structural, energetic, and experimental kinetic data were integrated into a structured database for model training and benchmarking.

We compared three predictive strategies: (1) Ligand-level MD, in which the overall 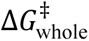 is directly converted into a *k*_off_ estimate; (2) Fragment-level MD, where individual fragment-specific 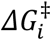 values are aggregated using a sum-of-barriers approach to approximate the total dissociation barrier; and (3) DCBK-ML, a machine-learning model integrating simulation-derived features with ligand and protein descriptors while retaining fragment-level interpretability. Within DCBK-ML, the cumulative feature stages comprised ligand molecular fingerprints (FP), complex-context descriptors (Cpx), ligand physicochemical descriptors (PC), the whole-ligand predicted activation free energy term (whole-ΔG^‡^), and the final fragment-informed block containing both fragment ΔG^‡^ terms and fragment-SMILES-derived features (Frag).

Performance was quantified using Pearson correlation (Pearson r) and mean squared error (MSE) against experimental values. On the HSP90 benchmark, the final DCBK-ML ensemble, defined as the mean prediction of XGBR, RF, and ExtraTrees, achieved Pearson r = 0.90 and MSE = 0.21. This performance exceeded that of recent representative studies, including Zhou et al. (2023) (r = 0.73, MSE = 0.64), Zhao et al. (2023) (r = 0.87, MSE = 0.26), the mixture-of-experts kinetic model of Zhao et al. (2024) (r = 0.82, MSE = 0.30), and Su et al. (2020) (r = 0.62, MSE = 0.72), positioning DCBK-ML as a state-of-the-art framework for k_off_ prediction while retaining physically meaningful and interpretable descriptors (Fig. 2A, Table S2).

**Figure 2.**
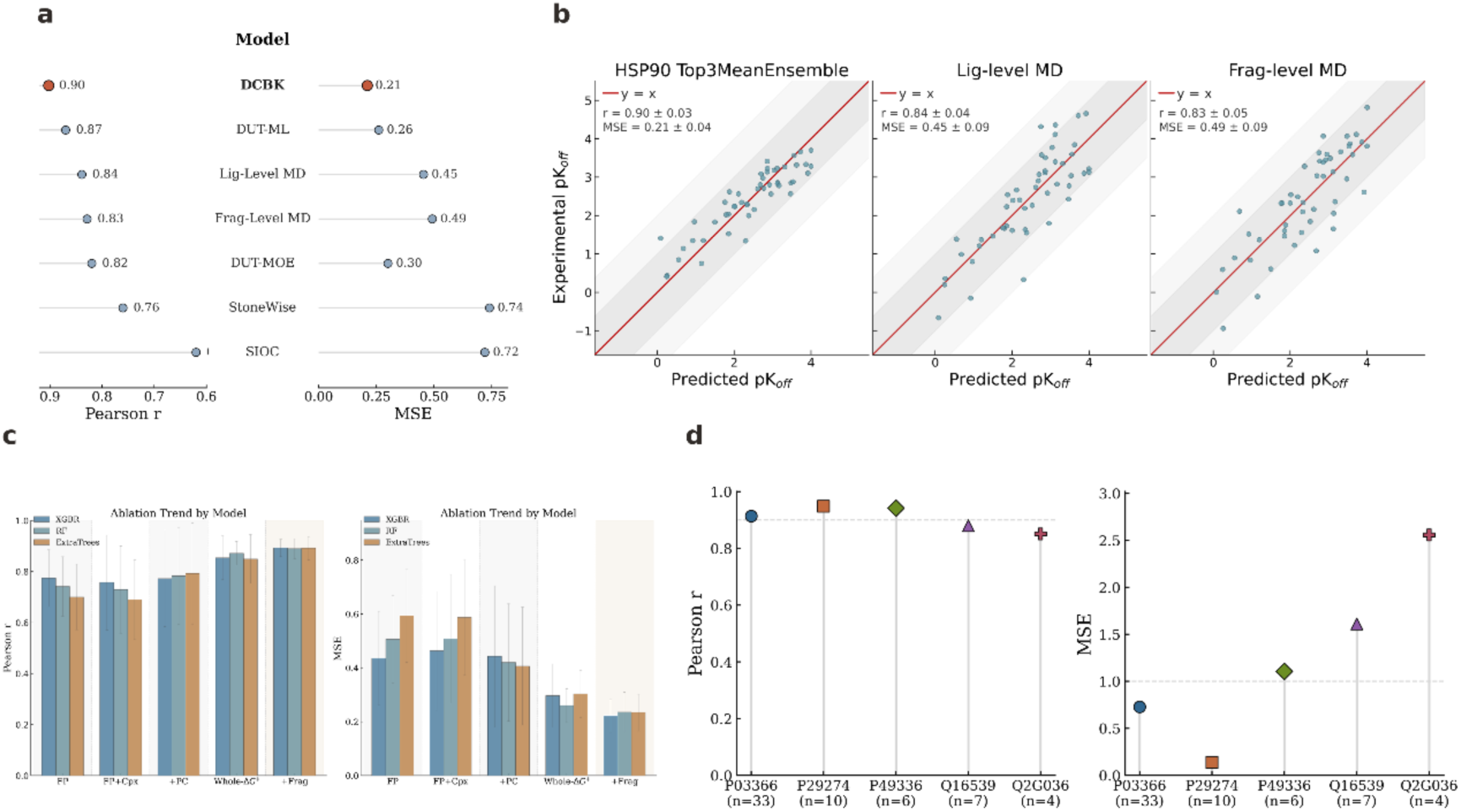
Performance of the hybrid simulation-ML framework across the HSP90 benchmark. (A) Quantitative comparison of DCBK-ML against representative recent studies using Pearson r and MSE, with the DCBK-ML row corresponding to the XGBR + RF + ExtraTrees mean ensemble. (B) Combined parity panel for the HSP90 ensemble model, ligand-level MD, and fragment-level MD representations. Within panel B, the red diagonal indicates ideal parity (y = x), the gray bands denote ±1 and ±2 units, and all subplots share the same axis range to make deviations directly comparable. (C) Cumulative HSP90 ablation analysis showing Pearson r and MSE (±STD) across the five feature stages FP, FP+Cpx, +PC, +Whole-ΔG^‡^, and +Frag for XGBR, RF, and ExtraTrees. (D) Cross-family transfer summary for the HSP90-trained XGBR + RF + ExtraTrees mean ensemble across the held-out protein systems.

In direct methodological comparisons, DCBK-ML also outperformed ligand-level MD (Pearson r = 0.84, MSE = 0.45) and fragment-level aggregation (Pearson r = 0.83, MSE = 0.49) across the full dataset (Fig. 2B, Table S3). Although fragment-based calculations captured part of the kinetic signal, they did not account for inter-fragment coupling; the hybrid ML layer systematically corrected these limitations and improved generalization.

Ablation analysis further showed that predictive accuracy improved as ligand fingerprints were progressively augmented with complex-context descriptors, ligand physicochemical properties, the whole-ligand predicted ΔG^‡^ term, and finally fragment-informed features. The final fragment-informed stage (+Frag) yielded the strongest HSP90 performance (Pearson r = 0.89, MSE = 0.22), indicating that fragment-resolved information is most effective when integrated with global ligand topology, physicochemical, and complex-context features (Fig. 2C).

Cross-family generalization of the HSP90-trained ensemble is summarized in Fig. 2D. Under held-out-family evaluation, the model retained transferable kinetic signal but showed system-dependent degradation, indicating that domain shift across protein families remains an important source of error beyond the core HSP90 benchmark. Among the optimized individual learners, XGB provided the strongest 5-fold performance on the 50-ligand HSP90 set (Pearson r = 0.90, MSE = 0.20), supporting its use as the leading single-model baseline within the ensemble framework.

### Cross-system validation and kinetic-regime analysis

We next evaluated cross-system generalization using leave-one-system-out (LOSO) validation across all six protein families (Fig. 3A-D). Among the five cumulative feature stages, the +Frag stage gave the strongest overall performance, with RF achieving a mean system-level Pearson r of 0.92 and a mean system-level MSE of 0.49; the corresponding pooled LOSO predictions reached Pearson r = 0.894 and MSE = 0.34. This represents a substantial improvement over the FP stage, for which the best pooled MSE remained 2.99. The progression across feature stages further showed that the major performance gain emerged after inclusion of the whole-ligand energetic prior: pooled Pearson r remained - 0.17 at the +PC stage, increased to 0.81 at +Whole-ΔG^‡^ stage, and improved further at +Frag stage. Thus, the LOSO analysis recapitulated the same cumulative trend observed in the HSP90-focused ablation, namely that global energetic information provides the dominant predictive foundation, while fragment-resolved features further refine cross-system transferability.

**Figure 3.**
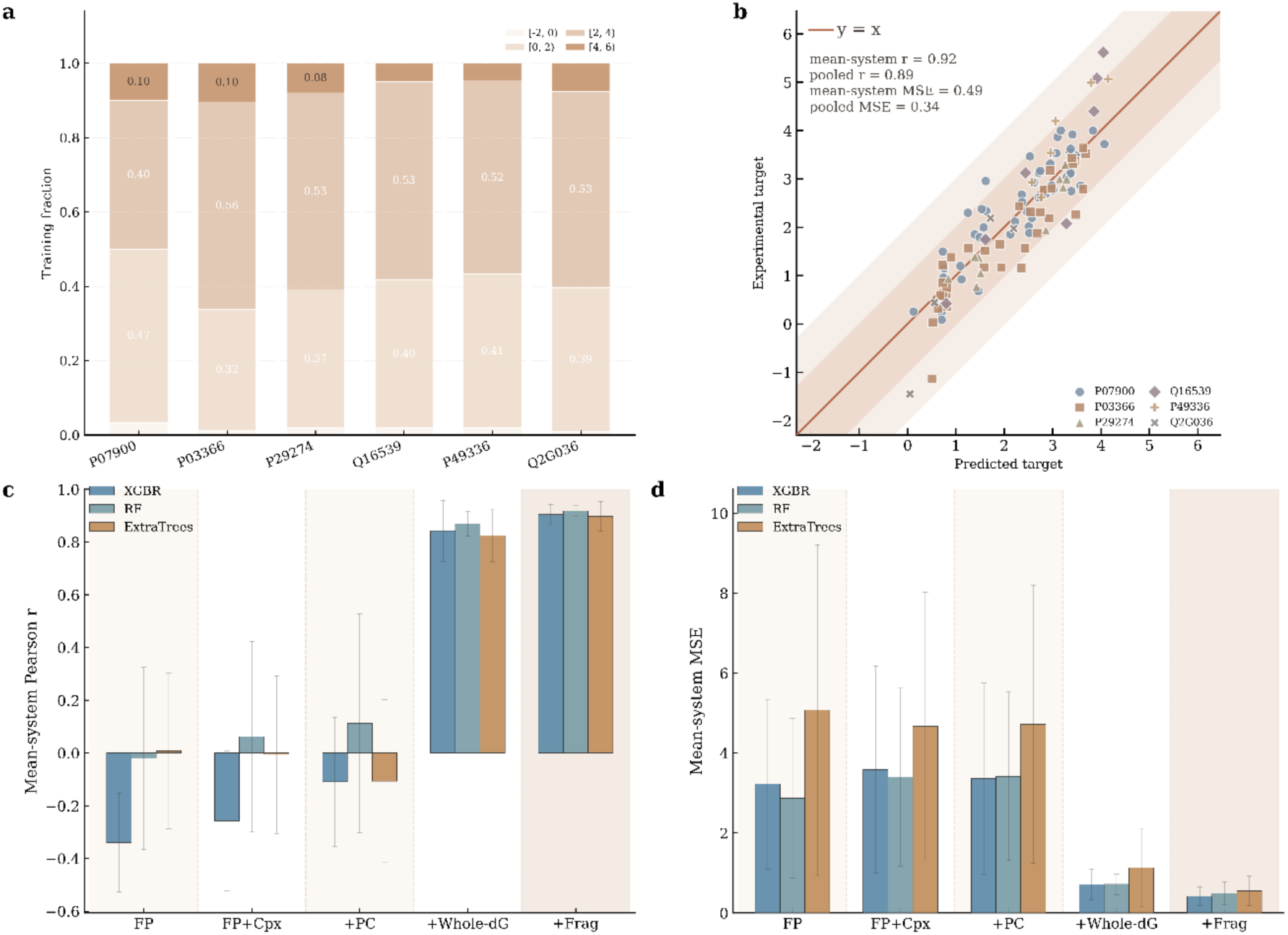
Leave-one-system-out validation and target-regime diagnostics. Panel A shows the training-target composition of each LOSO split. Panel B shows the pooled parity relationship between predicted and experimental targets for the best LOSO configuration, with points colored by protein system. Panels C and D report the mean system-level Pearson r and mean system-level MSE, respectively, for XGBR, RF, and ExtraTrees across the five cumulative feature stages from FP to +Frag stage.

System-level performance nevertheless remained heterogeneous across held-out families. The model performed best on the A_2A_R, for which the LOSO Pearson r reached 0.94 with an MSE of 0.18, whereas MAPK14 and CDK8 were the most challenging systems, with MSE values of 0.89 and 0.68, respectively. This family-dependent variation indicates that residual prediction error is not uniform across targets and is influenced by how well a held-out system is represented by the training distribution. Consistent with this interpretation, the target-composition map in Fig. 3A shows that each LOSO split is dominated by samples in the [0, 2) and [2, 4) target ranges, whereas high-target complexes are comparatively sparse across all splits.

Supplementary Fig. S1 provides additional diagnostic context for these cross-system results. The held-out-system error profile and distribution-shift analysis show that CDK8 exhibits the largest train-test mismatch, with a JS divergence of 0.37 and an absolute mean-target shift of 1.74, and it correspondingly ranks among the most difficult LOSO cases. Together, these results indicate that distribution mismatch, rather than sample count alone, is a principal constraint on cross-system generalization. Supplementary Fig. S1 also highlights the comparative parity behavior of the HSP90 Top3 ensemble, ligand-level MD, and fragment-level MD representations on a shared coordinate scale, allowing deviations from the ideal diagonal to be assessed consistently across methods. In parallel, the cumulative feature definitions used throughout the LOSO analysis remain aligned with the HSP90 ablation scheme, namely FP (ligand molecular fingerprints), Cpx (complex-context descriptors), PC (ligand physicochemical descriptors), Whole-ΔG^‡^ (the whole-ligand predicted ΔG^‡^ term), and Frag (the final fragment-informed block containing both fragment ΔG^‡^ terms and fragment-SMILES-derived features).

### Fragment-Level Insights and Interpretability

A key advantage of our framework is that it yields interpretable fragment-level signals that can be translated into medicinal-chemistry hypotheses. In HSP90, the matched analog pair corresponding to dataset indices 512→511 (Table S1) provides the clearest fragment-resolved success case. The two ligands differ only at the substituent vector assigned to fragment 2, and this localized modification is accompanied by an increase in pk_off_ from 1.85 to 2.19 (Δpk_off_ = +0.340). Consistently, the SHAP share associated with this fragment-2-defined region rises from 0.061 to 0.075, whereas the contributions from the other fragment-defined regions remain smaller. In this case, the model therefore points to a chemically specific optimization handle: strengthening the local fragment-2 substituent environment is associated with a slower dissociation phenotype (Fig. 4D).

**Figure 4.**
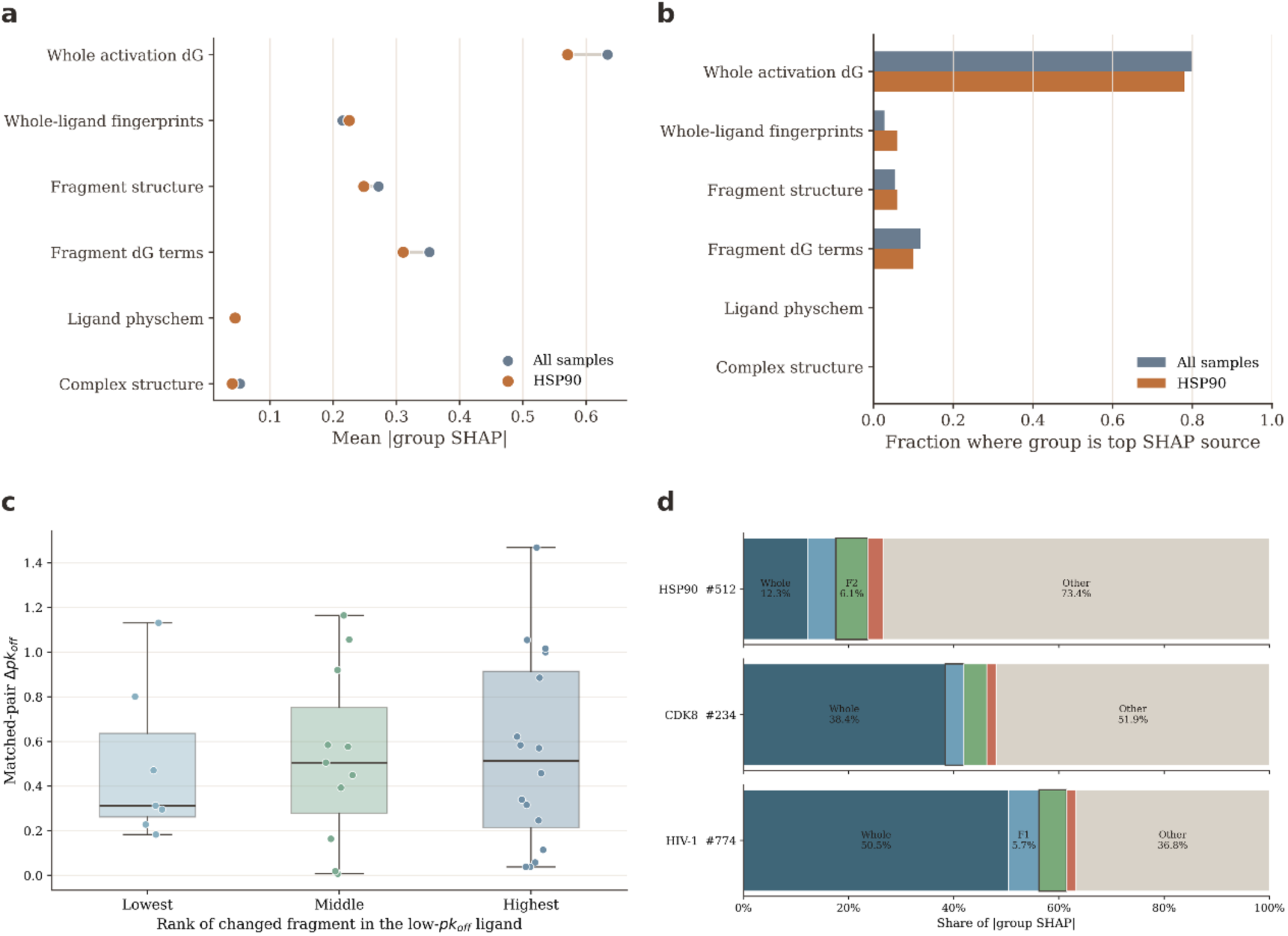
Global SHAP behavior and fragment-level interpretability across systems. Panel A shows the group-level SHAP importance for all samples versus the HSP90 subset. Panel B shows the fraction of samples in which each group is the top SHAP source for all samples versus HSP90. Panel C shows the cross-system matched-analog analysis using the full matched-pair set after excluding one visually dominant outlier from the middle-rank group. Panel D shows the grouped SHAP decomposition (whole, f1, f2, f3, and other) for the representative low-pk_off_ ligands from HSP90 (512), CDK8 (234), and HIV-1 (774); these shares identify the dominant model block, but the chemically actionable interpretation still requires inspection of the matched fragment structures.

In CDK8, the only clean matched analog pair is given by dataset indices 234→233 (Table S1). Here, the structural change is restricted to the region assigned to fragment 1, and pk_off_ increases from 2.618 to 3.538 (Δpk_off_ = +0.920). This trend is directionally consistent with a fragment-1-led optimization hypothesis. However, the SHAP contribution attributed to the modified fragment-1 region increases only modestly and remains smaller than the combined whole-ligand-plus-other signal. We therefore interpret CDK8 as a partial-support case: the fragment-1-defined substituent vector is the most plausible local handle for optimization, but the kinetic gain cannot be explained by a purely fragment-additive mechanism alone (Fig. 4D).

By contrast, HIV-1 is better represented by the matched pair corresponding to dataset indices 774→775 (Table S1), which exemplifies a more globally coupled regime. The chemical modification maps to the fragment-2-defined region, and pk_off_ increases from 0.580 to 1.164 (Δpk_off_ = +0.584), but the SHAP contribution assigned to that local change does not become dominant. Instead, the improved ligand is driven primarily by a redistribution toward the whole-ligand and other feature groups, indicating that the kinetic shift cannot be attributed to a single substituent change in isolation. HIV-1 therefore provides the clearest empirical example of the limits of a simple fragment-by-fragment interpretation and suggests that, in some systems, dissociation kinetics are governed by more collective structural effects along the unbinding pathway (Fig. 4D).

Together, these three systems delineate the principal interpretability regimes captured by DCBK-ML. HSP90 represents a case in which a localized substituent change is closely aligned with the kinetic improvement and is mirrored by a corresponding increase in the SHAP contribution of the same fragment-defined region. CDK8 represents an intermediate regime, in which one chemically modified region remains the most plausible local optimization handle, but the overall prediction is still shaped by broader whole-ligand and contextual effects. By contrast, HIV-1 exemplifies a more distributed regime, in which the kinetic gain cannot be assigned to a single substituent vector and instead reflects a broader redistribution of importance toward whole-ligand and residual feature blocks. Figure 4C summarizes this matched-analog analysis across systems using the full set of single-fragment analog pairs after exclusion of only the most extreme Δpk_off_ outlier from the middle-rank group. Figure 4D then shows the corresponding grouped SHAP decompositions for the representative lower-pk_off_ starting ligands with dataset indices 512, 234, and 774 (Table S1). Importantly, these grouped SHAP profiles identify the model component most strongly associated with the predicted kinetic behavior, but medicinal-chemistry interpretation still requires direct inspection of the underlying structural changes in the matched analog pairs. In this sense, the model does not replace chemical judgment; rather, it helps prioritize which local substitution pattern or broader ligand context is most likely to warrant optimization.

To assess model behavior across different kinetic regimes, we repeated 5-fold cross-validation after restricting the dataset to the top 100%, 75%, 50%, and 25% of complexes ranked by experimental pkoff (Fig. S2). Across these subsets (n = 110, 83, 55, and 28), pooled Pearson r values were 0.90, 0.85, 0.74, and 0.73, whereas the corresponding MSE values were 0.33, 0.26, 0.26, and 0.24. Thus, although correlation decreases as the analysis becomes restricted to the highest-pk_off_ complexes, absolute error remains relatively stable, indicating that the fragment-informed model retains meaningful predictive fidelity even in the more selective slow-dissociation regime.

Overall, the fragment-level analyses highlight how DCBK-ML achieves complementary layers of interpretability. The fragment decomposition offers mechanistic clarity by pinpointing rate-limiting interactions, whereas the ML model quantifies the relative importance of physicochemical and structural descriptors. Together, these insights bridge physics-based understanding and statistical inference, providing a transparent alternative to black-box or fully empirical approaches.

### Workflow Efficiency and Comparative Assessment

A central objective of our hybrid DCBK-ML framework is to reduce dependence on brute-force long-timescale MD while retaining physically grounded interpretability. In the current implementation, generating one training example required both fragment-level and whole-ligand simulations: the three fragment branches were run in parallel and required approximately 30 GPU-hours in total on NVIDIA V100 GPUs, whereas the corresponding whole-ligand umbrella-sampling workflow required approximately 35 GPU-hours at the sampling depth used here. The present model therefore still depends on physics-based data generation during model construction, but once trained, inference for a new ligand is rapid.

Alternative enhanced-sampling approaches offer different speed-accuracy trade-offs. τ-RAMD can provide useful residence-time ranking information on comparable timescales, but it is typically used after target-specific calibration and is most naturally interpreted in relative rather than absolute terms [19, 20]. WT-MetaD and APR can provide rich mechanistic detail, yet their total cost depends strongly on collective-variable choice, pathway complexity, and convergence requirements [21, 22]. Pure ML methods are essentially negligible at inference time [23], although they do not by themselves provide barrier-resolved physical interpretation.

The DCBK-ML pipeline is highly parallelizable: each fragment branch and each umbrella window can be evaluated independently, while ML inference for new compounds is effectively instantaneous once the model has been trained. In practice, this makes the workflow suitable for prioritization rather than exhaustive brute-force simulation, because one can first score many hypothetical analogs and then reserve the most expensive physics-based calculations for the most promising candidates.

We therefore view DCBK-ML not as a universal replacement for all enhanced-sampling workflows, but as a practical compromise between purely simulation-based kinetics estimation and purely data-driven prediction. The fragment and whole-ligand MD calculations preserve mechanistic grounding, whereas the supervised model amortizes that information across chemically related ligands and across targets represented in the training set.

Any wall-clock comparison should be interpreted cautiously because absolute cost depends on hardware, implementation details, system size, and the chosen convergence criteria. Nevertheless, the qualitative trade-off is consistent: methods that retain more explicit physical detail usually demand more simulation time, whereas purely statistical predictors are faster but less mechanistically transparent.

Overall, the DCBK-ML workflow balances accuracy, interpretability, and efficiency by combining reusable physics-based descriptors with rapid supervised inference. This balance makes the framework suitable both for cross-system kinetic prediction and for mechanism-informed ligand optimization in practical drug-discovery settings.

**Table 1.** Summary of representative computational methods for predicting ligand dissociation kinetics. The table lists the study, computational approach, target systems and ligand counts, reported computational cost, quantitative accuracy metrics, and key insights or notable features. Computational cost is reported where available, including wall times, simulation speed, or relative speedup over conventional MD. Accuracy metrics include correlation coefficients, MSE, mean absolute error, or qualitative assessments. Key insights highlight mechanistic interpretability, scalability, automation, and cross-family applicability. References correspond to the numbered studies cited in the main text.

| Method | System(s) | Count | Cost | Accuracy metrics | Reference |
| --- | --- | --- | --- | --- | --- |
| τRAMD + ML<br>(IFP analysis) | HSP90,<br>M2 GPCR | 4 | 0.5–2<br>days/GPU | Correlation with<br>experimental<br>residence times | [20] |
| SILCS + ML<br>(decision tree/NN) | 13 proteins | 329 | Not<br>quantified | Robustness claimed | [24] |
| MD/BD/milestoning<br>(SEEKR) | β-<br>cyclodextrin | 7 | 3.8 μs/ligand | High rank correlation<br>for $k_{\text{off}}$; effective $\Delta G$ | [27] |
| Infrequent<br>metadynamics + ML | μ-opioid<br>receptor<br>(GPCR) | 2 | Not reported | Predicted $k_{\text{off}}$ within<br>1 order of magnitude | [26] |
| τRAMD + ML<br>regression/classification | HSP90 | 94 | Not reported | $R^2 = 0.75$; MAE =<br>0.39 | [19] |
| PMRRP + PLS +<br>RAMD | HSP90 | 501 | Not reported | Pearson $r = 0.76$;<br>MSE = 0.32 | [25] |
| String simulations +<br>ML<br>(NN/GBDT) | CDK2 | 3 | Not reported | ML accuracy 79.5–<br>83.6%; $\Delta G$ barrier<br>~10 kcal/mol lower | [28] |
| RAVE (deep learning)<br>+ MD | Lysozyme<br>L99A | 1 | 7 orders<br>faster than<br>MD | Predicted $\Delta G$ within<br>2–3 kcal/mol | [23] |
| Infrequent<br>metadynamics | FKBP | 2 | 6–51x faster<br>than MD | Matches MD free<br>energy, pathway,<br>residence time | [21] |
| REVO/WExplore | Host–guest | 3 | Not reported | Predicted $\Delta G$ 4.2<br>kcal/mol lower | [22] |

## Discussion

The results presented here position DCBK-ML as a hybrid framework that links physically grounded dissociation descriptors to cross-system kinetic prediction with a degree of interpretability that is difficult to obtain from either simulations or machine learning alone. On the HSP90 benchmark, the ensemble model outperformed both the direct whole-ligand MD baseline and the summed-fragment MD baseline, and the same cumulative feature logic was preserved under leave-one-system-out validation across six protein systems. Two conclusions emerge most clearly from these analyses. First, the whole-ligand predicted ΔG^‡^ term provides the dominant predictive foundation: the largest performance gain appeared when this global barrier descriptor was introduced into the feature space. Second, fragment-resolved features nonetheless contributed additional predictive value, indicating that local substructural information carries non-redundant kinetic signal beyond the global barrier alone.

The fragment-level analyses further clarify how these signals should be interpreted in a medicinal-chemistry setting. In HSP90, a localized substituent change tracked both the experimental pk_off_ improvement and the redistribution of SHAP attribution, yielding the clearest example of a chemically actionable local optimization handle. CDK8 occupied a more intermediate regime, in which one modified region remained the most plausible driver of the kinetic gain, but the final prediction was still shaped by broader whole-ligand and contextual contributions. HIV-1 illustrated the complementary limit: the kinetic improvement could not be assigned to a single substituent vector and instead reflected a more distributed reweighting of model evidence across whole-ligand and residual feature blocks. DCBK-ML is therefore most useful not as a rigid fragment-ranking rule, but as a framework for distinguishing when a local substitution hypothesis is likely to be informative and when a broader scaffold- or context-level redesign may be required.

This balance between mechanistic grounding and statistical generalization defines the practical niche of the method. Fully simulation-based approaches can provide rich pathway-level information, but they are difficult to deploy systematically across large analog series, whereas purely data-driven predictors are fast but often provide limited physical guidance for optimization. DCBK-ML occupies a productive middle ground: the whole-ligand and fragment-level MD calculations retain an interpretable energetic basis, and the supervised layer amortizes that information across chemically related ligands and across targets represented in the training set. Encouragingly, the slow-binder subset analysis showed that absolute error remained comparatively stable even as the evaluation was restricted to progressively higher-pk_off_ complexes, suggesting that the fragment-informed representation retains useful resolution in the kinetic regime that is often most relevant for residence-time optimization.

At the same time, the present study also defines the current limits of the framework. The workflow relies on a one-dimensional dissociation coordinate and on BRICS-based ligand partitioning; both choices improve tractability, but they may miss alternative exit pathways, coupled motions, or fragmentation schemes that are more chemically intuitive for a particular series. The dataset, although large by the standards of experimentally annotated dissociation kinetics, remains modest by machine-learning standards and does not yet cover covalent inhibitors, metalloproteins, or strongly pose-heterogeneous systems. In addition, grouped SHAP analysis identifies dominant model blocks rather than causal microscopic determinants; medicinal-chemistry conclusions still require direct inspection of the corresponding matched analog structures and, ideally, experimental validation. Future work should therefore focus on expanding target coverage, incorporating richer pathway representations and pose ensembles, and integrating more structure-aware learning architectures while preserving the mechanistic interpretability that is the central advantage of the DCBK strategy.

## Methods

### Complex Preparation and Parameterization

Protein-ligand complexes were assembled from the archived DCBK kinetic dataset, which consolidates PDBbind-derived structural entries together with literature-curated A_2A_R kinetics to broaden target coverage. The final curated table used for the main manuscript comprised 110 protein-ligand entries spanning HSP90 (n = 50), HIV-1 protease (n = 33), A_2A_R (n = 10), MAPK14 (n = 7), CDK8 (n = 6), and ClpP (n = 4). Only chemically complete ligands associated with experimental dissociation measurements were retained. High-quality X-ray structures were prioritized, and entries with unresolved binding-site chemistry that could not be repaired reliably were excluded from the final set. Covalent ligands and metalloprotein systems were not considered in the present workflow.

All complexes were prepared with AmberTools24. Protein coordinates were processed with pdb4amber to standardize residue naming, remove alternate conformers, and add missing heavy atoms where possible. Protonation states were assigned with PROPKA at pH 7.0. Missing loops or side chains close to the binding pocket were rebuilt only when the local binding geometry could be preserved; otherwise the entry was removed during curation. Proteins were parameterized with the AMBER ff14SB force field, whereas ligands were parameterized with GAFF2 using Antechamber-derived atom types and partial charges. Systems were solvated in explicit TIP3P water and neutralized with Na+ and Cl- to approximately 0.15 M ionic strength.

Each complex was embedded in an octahedral periodic box with at least 12 Å of solvent padding. Energy minimization comprised 10,000 steps, followed by heating from 0 to 300 K over 1 ns under NVT conditions with 50 kcal mol^−1^ Å^−2^ restraints on protein backbone and ligand heavy atoms. The systems were then equilibrated for 5 ns under NPT conditions at 300 K and 1 atm with progressive restraint release. Langevin dynamics was used for temperature control. These preparation steps produced relaxed starting structures for the subsequent biased dissociation simulations.

Preparation, file handling, and simulation setup were automated through the DCBK workflow scripts to ensure consistent coordinate processing, parameter assignment, and random-seed control across systems.

### Steered Molecular Dynamics for Pathway Generation

For each complex, an initial dissociation pathway was generated by steered molecular dynamics (SMD). The reaction coordinate was defined as the distance between the center of mass of three ligand atoms and three protein pocket atoms chosen from the closest bound-state contacts, providing a compact descriptor of ligand displacement away from the binding site. A moving harmonic restraint with a spring constant of 2.5 kcal mol^−1^ Å^−2^ was applied along this coordinate at a pull rate of 1 Å ns−1 until the ligand reached approximately 35 Å from the pocket reference.

Ligand-pocket contact counts, minimum protein-ligand distance, and cumulative work were monitored throughout the pulling process to ensure that the trajectory reflected a physically plausible release pathway rather than an anomalous distortion. When a trajectory failed these checks, an additional SMD replica with a different initial velocity seed was generated. Representative snapshots sampled along accepted SMD paths were used to initialize the umbrella-sampling windows.

### Umbrella windows and sampling

Umbrella-sampling (US) windows were placed along the SMD-derived reaction coordinate from the bound basin to the dissociated state, typically using approximately 30 windows with ∼1 Å spacing. Each window employed a harmonic restraint of 5 kcal mol^−1^ Å^−2^ centered at the target coordinate value. Starting structures for each window were minimized and briefly equilibrated before production sampling under NPT conditions at 300 K and 1 atm.

Production US trajectories were run for 5 ns per window. Potentials of mean force were reconstructed with WHAM using 0.1 Å bins, and uncertainties were estimated by 200 bootstrap resamplings. The activation barrier ΔG^‡^ was defined as the difference between the minimum of the bound-state basin and the highest point along the dissociation barrier on the reconstructed PMF. The same PMF-analysis pipeline was later applied to the fragment-level calculations, ensuring that whole-ligand and fragment-derived barrier terms were measured on a common operational scale.

### Fragmentation of Ligands and Fragment-Level US

To enable fragment-resolved kinetic analysis, ligands were decomposed into chemically meaningful substructures using RDKit BRICS fragmentation. BRICS cut sites were capped with hydrogens to yield chemically valid fragment molecules, and the resulting fragments were checked for sensible valence states. In most cases, the final decomposition produced three fragment branches, which were used consistently throughout the fragment-level simulations and SHAP analyses.

Each fragment was parameterized with the same ligand force-field protocol as the parent molecule. Fragment-level SMD and umbrella sampling were then carried out along the same dissociation coordinate defined for the intact complex so that fragment-specific barrier estimates could be interpreted relative to a common unbinding geometry. This approximation does not imply that isolated fragments unbind independently in the physical system; rather, it provides a controlled way to quantify the local energetic contribution associated with each fragment-defined region of the parent ligand.

For the fragment-level MD baseline, the fragment-specific ΔG^‡^ terms were aggregated into a summed-barrier descriptor, and a calibration from computed barrier quantities to the experimental pk_off_ scale was estimated within training data only before being applied to held-out samples. In the DCBK-ML setting, the individual fragment ΔG^‡^ values were retained as separate model inputs in addition to this aggregate view, enabling the hybrid framework to capture both local and global kinetic contributions.

### ML Model Integration

The machine-learning analyses were built on the curated table of 110 protein-ligand entries. The experimental target used throughout the manuscript was pk_off_, defined on the stored logarithmic kinetic scale of the curated dataset. In the global leave-one-system-out analyses, this stored target was used directly without applying a second transformation. Dataset indices cited in the main text and in Table S1 correspond directly to the index column of this curated table.

Input features were organized into the same five cumulative stages used in the Results section: FP, FP+Cpx, +PC, +Whole-ΔG^‡^, and +Frag. FP comprised a 200-bit Morgan fingerprint of the full ligand. FP+Cpx added latent complex-structure components derived from contact- and SASA-based descriptors when a usable complex structure was available; these features were standardized, compressed by PCA, and the retained components were selected within the training partition by Lasso. Samples lacking structure-derived inputs were carried forward with missing values and handled by split-specific median imputation. +PC added eight ligand physicochemical descriptors: molecular weight, logP, hydrogen-bond donors, hydrogen-bond acceptors, topological polar surface area, rotatable bonds, aromatic ring count, and total ring count. +Whole-ΔG^‡^ introduced the whole-ligand predicted ΔG^‡^ term. +Frag added fragment-specific ΔG^‡^ values together with fragment-SMILES-derived descriptor and similarity features.

Model selection proceeded in two stages. An initial HSP90-specific screen compared the optimized five-model panel used in the Results section, after which the final benchmark was carried forward using the top three base learners: XGBR, RF, and ExtraTrees. For the HSP90-specific benchmark, the 50 entries with system label P07900 were evaluated under 5-fold cross-validation with split-specific median imputation and standardization, and the final DCBK-ML benchmark was reported as the mean prediction of these three models. The same three regressors were retrained across all cumulative feature stages for the HSP90 ablation analysis shown in Fig. 2.

For cross-system evaluation, leave-one-system-out validation was applied across the six protein systems. In each split, one system was held out entirely for testing, and all preprocessing steps were learned from the training partition only to prevent information leakage. Model performance was summarized using both pooled and system-level Pearson r and MSE. Additional analyses restricted the dataset to the top 100%, 75%, 50%, and 25% of entries ranked by experimental pk_off_ to assess behavior in progressively slower-dissociating regimes; this subset analysis was performed with the final fragment-informed RF configuration used for Supplementary Fig. S2.

Interpretability analyses were performed with grouped TreeSHAP values on the feature-complete XGBR model. For the global all-sample versus HSP90 comparison, features were aggregated into six blocks: whole-ligand activation, whole-ligand fingerprints, fragment structure, fragment dG terms, ligand physicochemical descriptors, and complex-structure components. For matched-analog and new-molecule analyses, the grouped output was reorganized into five medicinal-chemistry-oriented blocks: whole, f1-related, f2-related, f3-related, and other. The same grouped SHAP workflow can be applied to a new molecule once the corresponding whole-ligand and fragment-level descriptors have been generated.

### Data Management and Reproducibility

All curated kinetic measurements, simulation-derived barrier terms, and machine-learning features were stored in structured CSV tables. Table S1 provides the dataset-index mapping used throughout the manuscript, including the sample identifiers cited in the fragment-level matched-pair analyses. Each entry records the ligand identity, system annotation, experimental target value, and associated whole-ligand and fragment-level descriptors.

Simulation setup, SMD, umbrella sampling, PMF reconstruction, feature extraction, model training, SHAP analysis, and figure generation were automated through version-controlled Python and shell scripts. Reproducibility packages accompanying the main and supplementary figures include the processed inputs, plotting scripts, summary tables, and output artifacts required to regenerate the reported analyses.

Software versions for AmberTools, RDKit, scikit-learn, XGBoost, and related analysis dependencies were recorded within the distributed workflow. Intermediate artifacts, including PMF curves, cross-validation predictions, and grouped SHAP summaries, were retained to facilitate independent verification and re-analysis.

## Data Availability

Data and Code Availability. All data supporting the findings of this study are provided within the paper and its Supplementary Information, including the curated dataset of protein-ligand complexes with experimental pk_off_ values, the computed whole-ligand and fragment-level ΔG^‡^ terms, and the machine-learning feature tables used for model training and evaluation. Table S1 provides the dataset-index mapping used throughout the main text.

Processed input files, analysis tables, figure-specific reproducibility packages, and the Python and shell scripts used for simulation setup, PMF analysis, feature generation, model training, SHAP analysis, and figure production are available from the corresponding author upon reasonable request. Publicly sourced structural and kinetic data, including entries originating from PDBbind 2020 and literature-curated measurements, were obtained from their original sources.

## Author Contributions

All authors contributed to the research and manuscript preparation. R.Z. and J.G. conceived the study and supervised the project. Y.L. developed the computational methodology and performed the molecular dynamics simulations. Y.W., X.Z., and J.G. implemented the ML model and carried out data analysis. R.Z. and Y.L. designed the fragment-based strategy. Y.L., Y.W., X.Z., and J.G. integrated the workflow and conducted the validation experiments. R.Z. provided domain expertise on kinetics and helped curate experimental data. All authors discussed the results and contributed to writing and revising the manuscript.

## Support information

**Figure S1.**
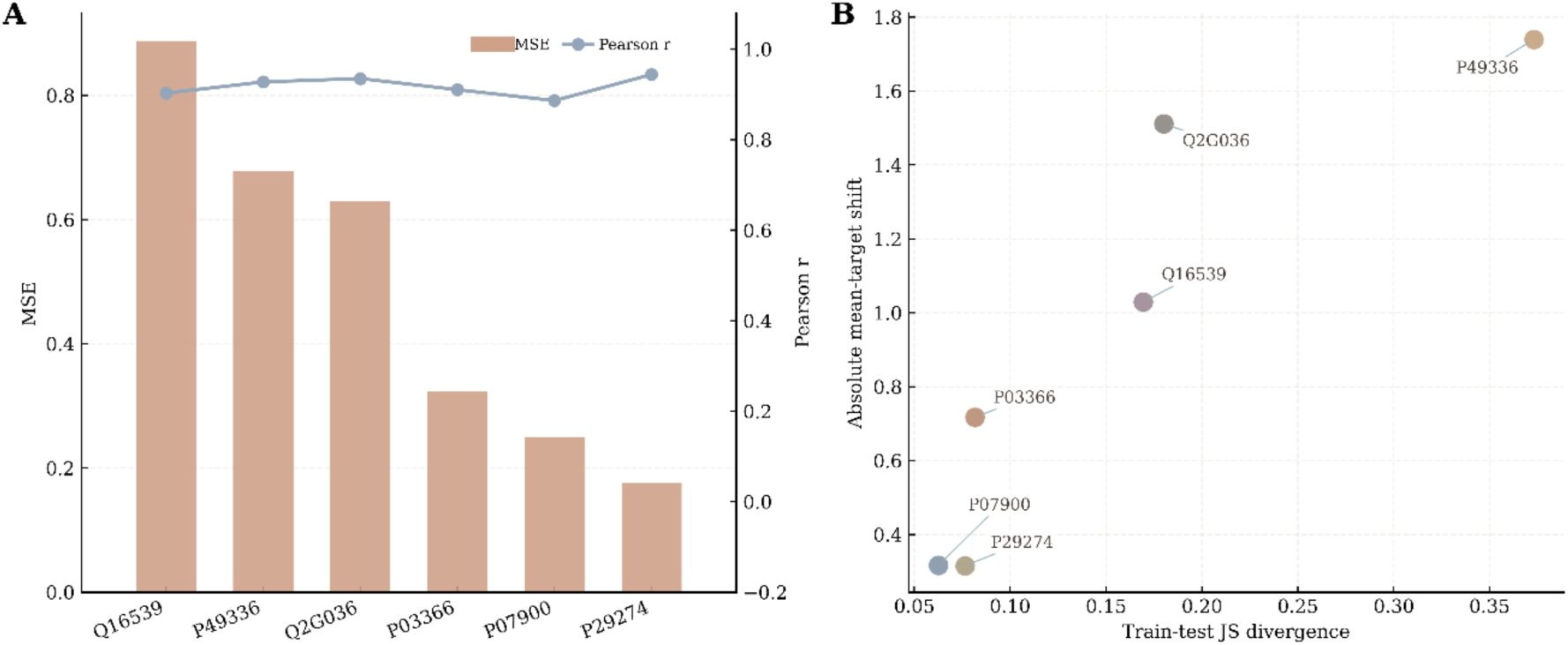
Supplementary LOSO diagnostics. Panel A summarizes the held-out-system error profile for the best energetic model under LOSO evaluation using MSE bars and Pearson r points. Panel B maps the corresponding train-test distribution shifts using JS divergence and absolute mean-target shift, highlighting systems with the largest domain mismatch.

**Figure S2.**
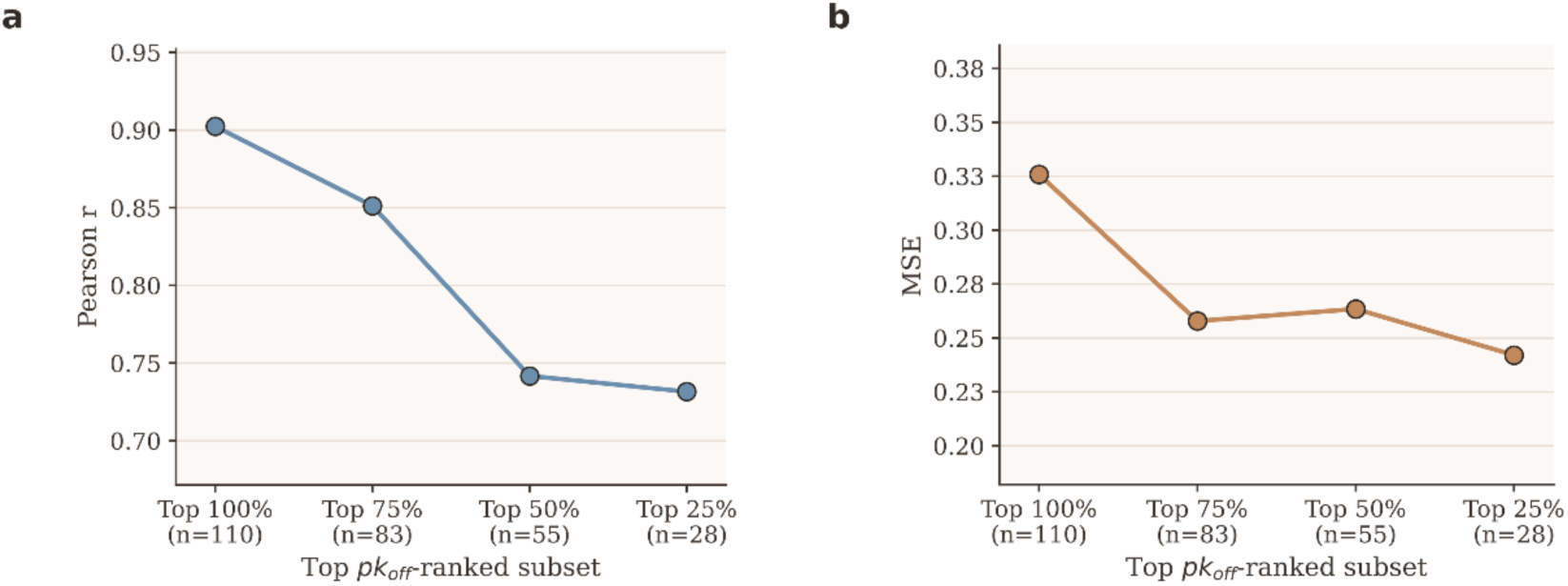
Top-pk_off_ subset performance of the final fragment-informed RF model. Panel A reports pooled 5-fold Pearson r for the top 100%, 75%, 50%, and 25% of complexes ranked by experimental pk_off_. Panel B reports the corresponding RMSE values for the same ranked subsets.

